# A proximity biotinylation approach for the identification of membrane contact site proteins in *Toxoplasma gondii*

**DOI:** 10.64898/2026.08.05.743015

**Authors:** Kaelynn V. Parker, Diego Huet

**Affiliations:** Department of Cellular Biology, University of Georgia, Athens, GA, USA; Department of Pharmaceutical and Biomedical Sciences, University of Georgia, Athens, GA, USA; Center for Tropical and Emerging Global Diseases, University of Georgia, Athens, GA, USA

**Keywords:** *Toxoplasma*, Organelle, Apicoplast, Mitochondrion, Endoplasmic reticulum, Membrane contact sites, Proximity biotinylation

## Abstract

Inter-organellar communication is crucial for cellular function. Inside the cell, organelles interact with each other via membrane contact sites (MCSs). These structures mediate the close apposition of two organellar membranes to allow for the exchange of metabolites, ions and lipids. Most of what is known about MCSs comes only from a handful of well-studied metazoans, particularly yeast and mammals. Apicomplexans are parasites that drive human disease throughout the world. Yet, little is known about the makeup or function of their MCSs, leaving a gap in our understanding of how organelles communicate beyond conventional model eukaryotes. Here, we used a proximity biotinylation approach to map the surface proteome of three organelles in the model apicomplexan *Toxoplasma gondii*: the apicoplast—a non-photosynthetic plastid found only in apicomplexans—its single mitochondrion and the endoplasmic reticulum. By subtracting a cytosolic spatial reference, our high-stringency proteomic analysis uncovered candidate proteins localized simultaneously to multiple organellar surfaces suggesting their role as MCS components. We then validate our approach by characterizing a candidate involved in the association between the apicoplast and the mitochondrion. Overall, our findings provide a valuable approach to identify MCSs in apicomplexans and set the stage to apply our approach to other organelles in these pathogens.

**Highlights:**

- Generation of surface proteomes for the apicoplast, mitochondrion, and ER in *Toxoplasma gondii*
- Mapping of the first endoplasmic reticulum and mitochondrial surface proteomes in *T. gondii*
- Identified novel membrane contact site candidate proteins
- Validated a membrane contact site candidate mediating mitochondrion–apicoplast interactions

## Introduction

Eukaryotic cells are defined by membrane-bound organelles that compartmentalize specific biochemical processes. To maintain homeostasis, organelles must communicate with one another, which can be achieved through vesicular trafficking, cytoplasmic diffusion or via specialized regions termed membrane contact sites (MCSs)^1^. MCSs are functional close contacts between organelles used to exchange proteins, ions, lipids and signaling molecules^2^. These areas also play crucial roles in autophagy and organelle division. Consequently, their dysregulation is implicated in various human diseases, highlighting their central role for cellular homeostasis^3,4^.

Apicomplexans are a massive phylum of parasitic protists. All species are obligatory parasites, and many cause devastating diseases in humans and animals. Apicomplexans in the genus *Plasmodium* cause malaria, which resulted in 282 million cases and 610,000 deaths in 2025^5^. *Toxoplasma gondii*, one of the most successful apicomplexans, chronically infects a third of the world’s population and can cause fetal abnormalities during pregnancy and opportunistic brain infections in immunocompromised patients^6–8^. Apicomplexans are also responsible for important livestock diseases, inflicting billions of dollars in annual economic losses^9,10^. Collectively, these parasites impose a massive global health and economic burden, with the highest toll falling on developing nations. Most apicomplexans harbor two organelles of endosymbiotic origin: a single mitochondrion and a relict, nonphotosynthetic plastid named the apicoplast. These two compartments have coevolved for at least 400 million years, forming intimate physical and biochemical connections that result in chimeric biosynthetic pathways shared between them^11–13^. For instance, isoprenoid and heme biosynthetic intermediates are exchanged between the apicoplast and the mitochondrion^14–16^. While all eukaryotic cells rely on inter-organellar communication, our knowledge of MCS structure and function stems almost entirely from studies in opisthokonts, a clade of eukaryotes that includes mammals and yeast, and plants^17,18^. Although contact sites have been reported in *Plasmodium* and *T. gondii*^19–22^, our knowledge remains fragmentary compared to well-studied model organisms ^23,24^. To identify MCS protein candidates, the interface between different organelles must be studied. This is a challenging endeavor, as our knowledge of *T. gondii* organellar surface proteins is still very limited. This gap is largely driven by the evolutionary divergence and specialization of apicomplexan organelles, as well as the fact that approximately half of their proteome consists of uncharacterized, phylum-specific “hypothetical” proteins.

Here, we have used proximity biotinylation for proteomic mapping of the outer membranes of the apicoplast, the mitochondrion and the endoplasmic reticulum of *T. gondii*. Together with a spatial reference control, this unbiased approach further allowed us to identify potential MCS candidates, and we validate our approach by characterizing a protein involved in the mitochondrion-apicoplast association.

Our work exemplifies the power of proximity labeling for studying organellar biology in apicomplexans and provides a methodological framework for interrogating the surface proteome of other subcellular compartments in this important group of pathogens.

## Results

### Generation of molecular handles to probe organellar interfaces

Because our knowledge of *T. gondii* organelle surface proteins remains limited, investigating organellar interfaces to identify MCSs presents a significant challenge. To overcome this hurdle, we chose the biotin ligase enzyme TurboID^25^ (TID) for proximity biotinylation. This labeling approach has been implemented for the proteomic mapping of the mammalian outer mitochondrial membrane^26^, as well as to identify endoplasmic reticulum (ER)-endosomes^27^, ER-plasma membrane contacts^28^and lipid droplet-mitochondria contacts^29^ in an unbiased manner. Before beginning our experiments, we sought to target GFP (as a proxy for TID) specifically to the outer membranes of 3 different organelles: the apicoplast, the mitochondrion and the ER.

To assess whether GFP could be targeted to the outer apicoplast membrane, we endogenously tagged the apicoplast triose phosphate translocator 1 (APT1) with GFP using CRISPR/Cas9 as an organelle-specific localization handle (**Supplemental Figure 1A**). Because both the N-and C-termini of APT1 extend into the cytosol^30^, fusing GFP to the C terminus of APT1 should position the fluorescent reporter at the apicoplast surface. Furthermore, APT1 constructs have previously proven effective as molecular handles for identifying apicoplast transporters^31^ and for purifying apicoplasts in *Plasmodium falciparum*^32^. For generating our mitochondrial handle, we used the mitochondrial outer membrane localization sequence of human OMP25^33^ (**Supplemental Figure 1B**), which has been employed to expose GFP to the outer mitochondrial membrane of *T. gondii*^34^. Finally, to target the cytosol-facing endoplasmic reticulum membrane, we adapted a strategy from human cell studies that uses the C-terminal localization signal of cytochrome b5^35^ (Cb5). We identified and utilized the targeting signal from the *T. gondii* Cb5 homolog (TGME49_276110) in a plasmid for our ER handle (**Figure 1A**). Fusing this signal to GFP confirmed the colocalization of the fluorescent protein with calumenin, an ER protein, by immunofluorescence assay (IFA) (**Figure 1B**). We next assessed the topology of our handles and confirmed that their GFP tags are anchored to their respective organelle outer membrane and exposed to the cytosol. To test this, we employed a live-cell imaging assay using a cytosolic anti-GFP/YFP single-chain nanobody fused to mCherry^34^ (**Figure 1C**). After transiently transfecting a plasmid enabling the expression of the mCherry nanobody in our organellar handle-expressing strains, the mCherry nanobody was recruited to the organelle surface by all our candidate handles, confirming that their GFP tags face the cytosol, as expected (**Figure 1D**). In contrast, the nanobody remained diffuse throughout the cytosol in the absence of a GFP tag or when GFP was sequestered inside the ER lumen (GFP-HDEL) (**Figure 1D**).

**Figure 1:**
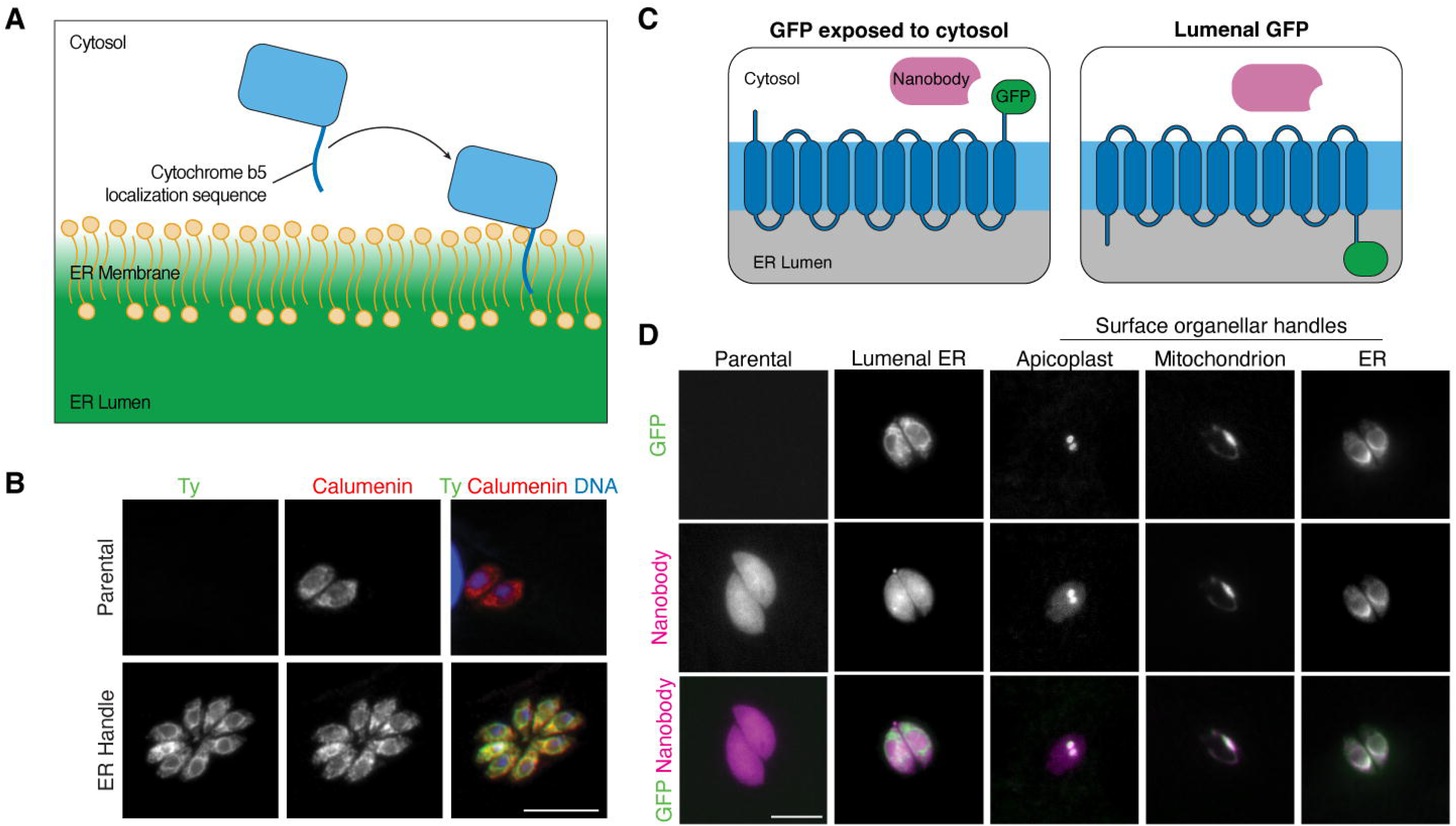
An ER-handle to target proteins to the surface of the ER in *T.gondii*. **(A)** Schematic of cytochrome b5 tail anchoring to ER membrane. **(B)** Intracellular parasites expressing the ER were fixed and stained for GFP (green), the ER marker calumenin (red), and DAPI (blue). **(C)** Schematic representation of the GFP-nanobody experiment to determine the topology of our organellar handles. **(D)** Live microscopy images showing the localization of the mCherry-tagged anti-GFP nanobody in strains expressing organelle-surface GFP handles or ER luminal GFP. As expected, the nanobody relocalizes to the cytosolic surface of the apicoplast, mitochondrion, and ER where the GFP epitope is exposed. In contrast, the nanobody does not interact with GFP localized within the ER lumen. Live imaging was performed 24 hours post transfection to visualize GFP (green) and GFP-nanobody (magenta). Scale bars: 5 µm.

### Characterizing activity of the organellar TurboID handles

Having confirmed the topology of our handles, we generated a new strain, termed AP-HA^TID^, in which we replaced GFP with TID in the APT1 locus to target the ligase to the apicoplast. We also generated two new organelle-targeted TID plasmids: MT-HA^TID^ (mitochondrion) and ER-Ty^TID^ (endoplasmic reticulum) (**Figure 2A and Supplemental Figure 2A-C**). In addition, we generated a strain where we endogenously tagged the C terminus of the non-essential cytosolic protein FBXO14^36^ with TID (**Figure 2A and Supplemental Figure 2D**). Termed CT-HA^TID^, this strain will serve as a spatial reference for our proteomic experiments, allowing us to distinguish proteins labeled specifically by our organellar handles from cytosolic bystanders. IFAs of parasites expressing our organellar handles confirmed the correct targeting of all four of them (**Figure 2B**). AP-HA^TID^ showed clear overlap with the apicoplast marker CPN60, while MT-HA^TID^, ER-Ty^TID^, and CT-HA^TID^ colocalized with TOM40 (mitochondrion), the ER-resident calcium pump SERCA (ER), and CDPK1 (cytosol), respectively. Cells expressing our handles treated with biotin for 4 hours and probed with an anti-streptavidin antibody show compartment-specific biotinylation (**Supplemental Figure 2E-F**), and western blot analysis of biotinylated proteins from whole cell lysates confirmed the activity of our handles (**Figure 2C and Supplemental Figure 2G**).

**Figure 2:**
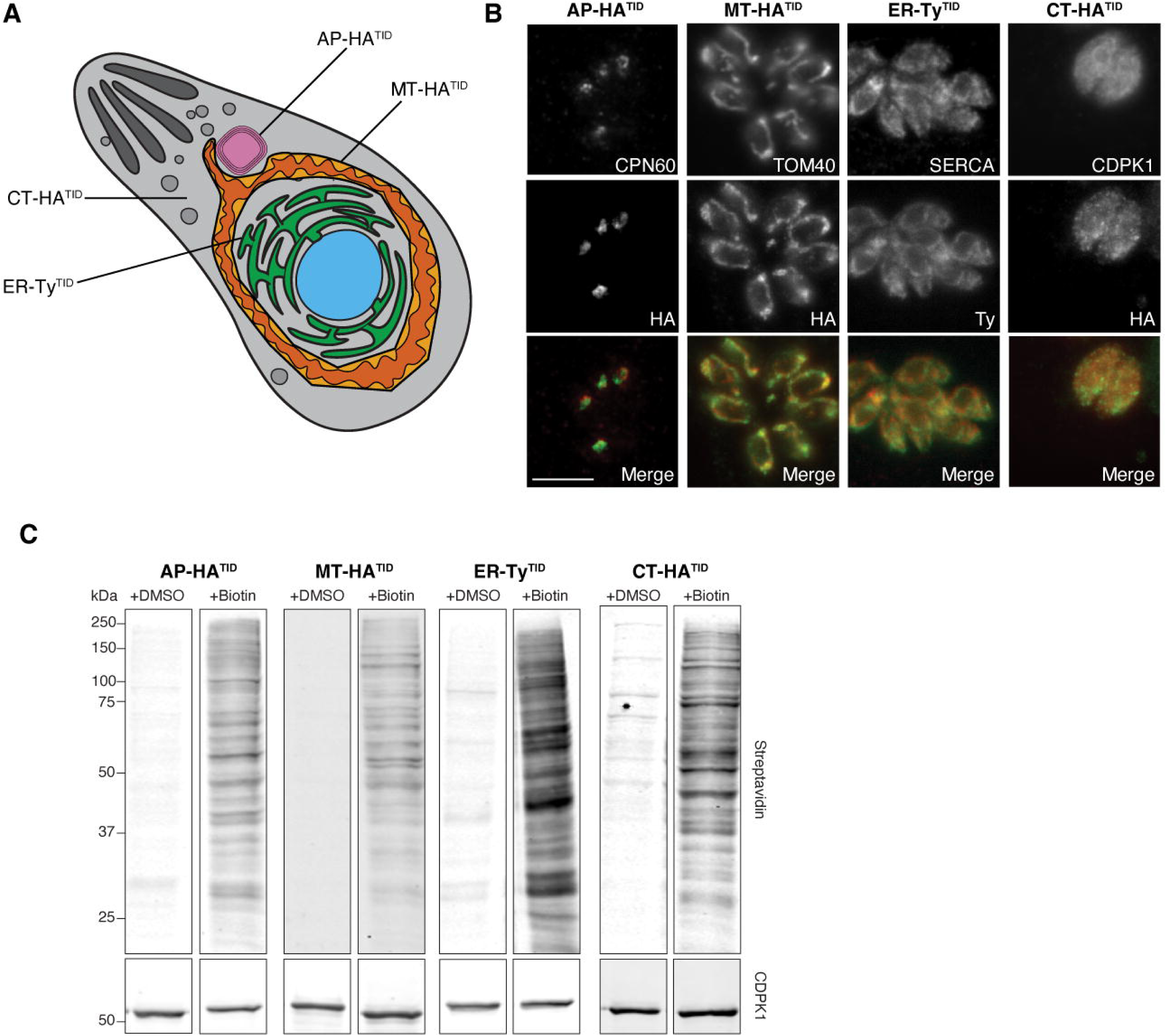
Organellar surface TurboID handles are correctly localized and enzymatically active. **(A)** Diagram illustrating the mitochondrion (orange), apicoplast (pink), and ER (green) in *T. gondii*. **(B)** The AP-HA^TID^, MT-HA^TID^, ER-TY^TID^ and CT-HA^TID^ strains were fixed and stained for the HA or Ty tags (green), alongside organelle-specific markers (red): CPN60 (apicoplast), TOM40 (mitochondrion), SERCA (ER), and CDPK1 (cytosol). Scale bar: 5 µm. **(C)** Parasites expressing AP-HA^TID^, MT-HA^TID^, ER-TY^TID^ or CT-HA^TID^ were incubated with 50 μM biotin or DMSO (vehicle control) for 4 hours and then lysed. Whole-cell lysates were analyzed by streptavidin blotting to detect biotinylated proteins (upper panels) and CDPK1 was used as a loading control (lower panels).

### Generation of organellar surface proteomes

After verifying the activity of our organelle-targeted TID handles, we incubated the parental line alongside the AP-HA^TID^, MT-HA^TID^, ER-Ty^TID^, and CT-HA^TID^ strains with biotin or DMSO (vehicle control) for 4 hours. Whole-cell lysates were incubated with streptavidin-coated magnetic beads to capture biotinylated proteins, and eluted fractions were analyzed by western blot to confirm successful enrichment of biotinylated proteins in experimental samples compared with negative controls (**Supplemental Figure 3A-B**).

Biotinylated proteins bound to the streptavidin beads were released by on-bead trypsin digestion and analyzed by liquid chromatography–tandem mass spectrometry (LC-MS/MS). Raw mass spectrometry data from three biological replicates per condition (incubated with DMSO or biotin) were processed for analysis, retaining only proteins detected with two or more unique peptides (**Figure 3A**). We also removed two highly expressed, endogenously biotinylated proteins from our analysis: the apicoplast-resident ACC1 (TGME49_221320)^37^ and PyC (TGME49_284190), a pyruvate carboxylase that localizes to the mitochondrial matrix^38^. To distinguish proteins specifically labeled by our organellar handles from non-specific cytosolic background, we applied a two-step filtering strategy: first, we calculated protein enrichment in biotin-treated samples relative to their respective negative controls; second, we identified proteins preferentially labeled by each organelle-targeted handle compared with the cytosolic reference strain, CT-HA^TID^, both in the presence of biotin (**Figure 3A**). Applying these filtering criteria yielded refined surface proteomes for the apicoplast (120 proteins), mitochondrion (40 proteins), and ER (44 proteins). Encouragingly, APT1 was identified among the top hits in our apicoplast dataset, validating our approach. Additionally, several of these proteins have established apicoplast localizations (**Figure 3B**, **Supplementary Table 1**). Amongst them, we found five hypothetical apicoplast proteins (HAPs 1, 11, 29, 45 and 46) previously localized to the organelle^31^; TgFLP12, a transporter localized to the apicoplast membranes^39^ and TgPL2, a protein found in the periphery of the apicoplast where it plays a crucial role in apicoplast integrity^40^. We also detected three autophagy-related proteins: TgATG8, known to localize to the outer membrane of the apicoplast^41^ as well as TgATG5 and TgATG16L, which reside as a complex at the apicoplast periphery^42^. Finally, we also found TgTPC, an apicoplast channel that mediates functional and physical coupling with the ER through inter-organellar calcium transfer^43^.

**Figure 3:**
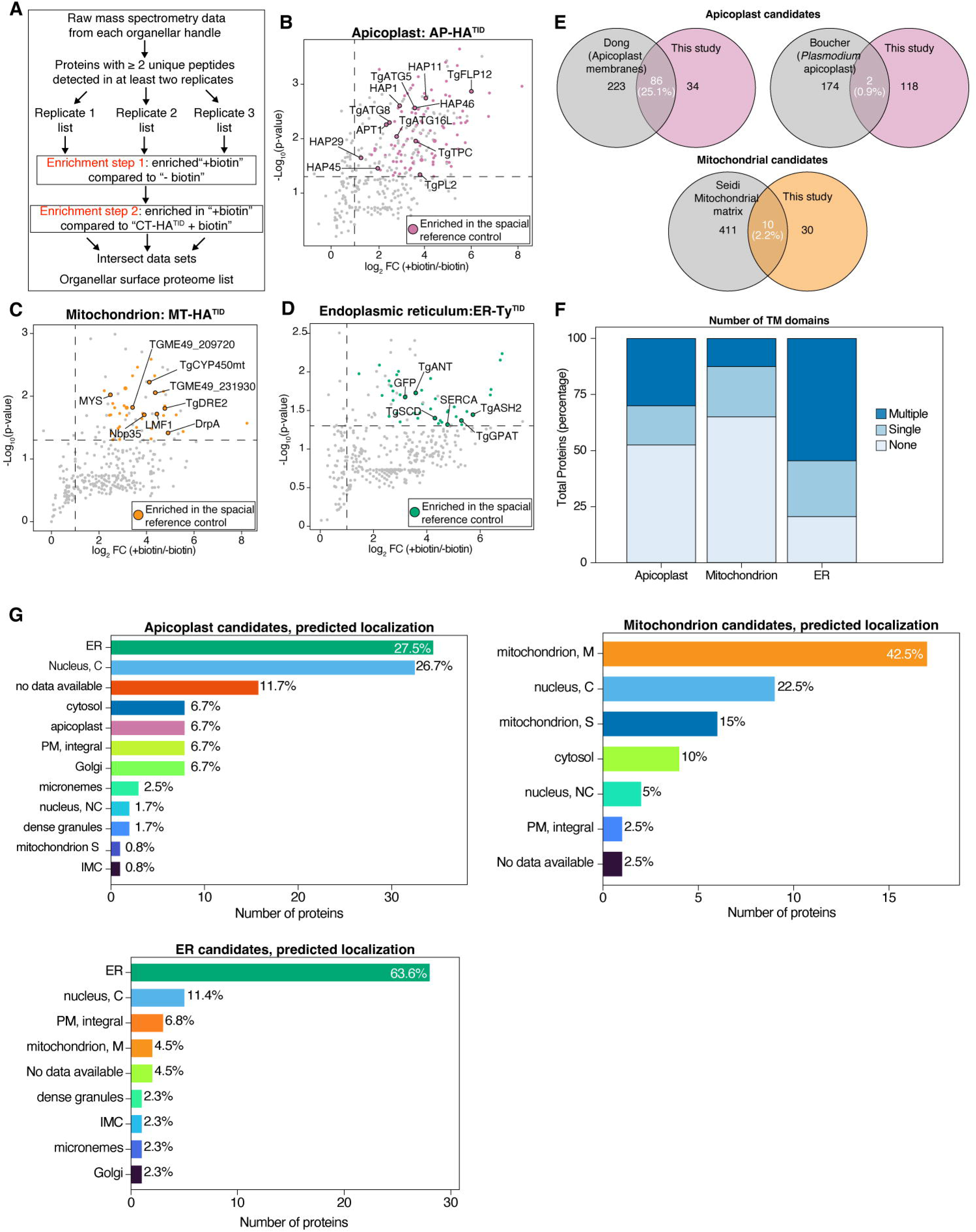
Proteomic mapping of the apicoplast, mitochondrion and ER surfaces. **(A)** Experimental design for mass spectrometry-based proteomics. Volcano plots showing enrichment of biotinylated proteins in the AP-HA^TID^ **(B)**, MT-HA^TID^ **(C)**, and ER-Ty^TID^ **(D)** strains in the presence versus absence of biotin. For each plot, the x-axis represents log_2_ fold in protein enrichment (+biotin / −biotin) and the y-axis represents the −Log_10_(p-value). Proteins significantly enriched in the spatial reference control (CT-HA^TID^) along with selected proteins with experimentally confirmed subcellular localizations from the literature are highlighted in each panel. **(E)** Venn diagrams showing the overlap between our apicoplast surface proteome (pink) and the apicoplast membrane and lumenal proteomes (grey), as well as our mitochondrial surface proteome (orange) versus the mitochondrial matrix proteome (grey). **(F)** Stacked bar chart showing the proportion of single-pass and multi-pass integral membrane proteins across our surface proteomes. **(G)** Predicted subcellular localizations mapped via hyper LOPIT of our identified apicoplast, mitochondrial, and ER surface candidates. Nucleus, C: Nucleus - Chromatin; Nucleus, NC: nucleus, non-chromatin; Mitochondrion M: Mitochondrion - membranes; Mitochondrion S: Mitochondrion - soluble; PM: plasma membrane.

In our mitochondrial surface dataset, we were unable to detect OMP25, likely due to its small size (30 amino acids) and the resulting limitations in tryptic peptide detection via mass spectrometry. However, we identified candidates that have been previously localized to the mitochondrion (**Figure 3C**, **Supplementary Table 1**). These include the outer mitochondrial membrane (OMM) MYS^44^, TGME49_231930^45^, TGME49_209720^46^ and LMF1^47^. Notably, LMF1 interacts with IMC10 to form MCSs between the mitochondrion and the inner membrane complex^21^. We also detected two components of the cytosolic iron-sulfur cluster assembly (CIA) pathway of *T. gondii*^48^: TgNbp35, which is anchored to the outer face of the OMM^49^, and TgDRE2, a predominantly cytosolic protein also found at the cytosol-mitochondrial interface in other organisms^50^. Similarly, we detected DrpA, which predominantly localizes to the apicoplast but has been observed colocalizing with the mitochondrion^51^.

In our ER surface data set, GFP—a component of our ER-TY^TID^ construct—was detected among our top hits (**Figure 3D, Supplementary Table 1**). We also confirmed that six of our top candidates are known ER-localized proteins, including SERCA^52^, the adenine nucleotide translocator TgANT^53^, and the active serine hydrolase 2, TgASH2^54^. In addition, we detected TgSCD, which colocalizes with the ER marker Der1^55^, and TgGPAT, found to localize to the perinuclear, ER-like compartment of the parasite^56^.

Next, we benchmarked our expanded proteomic datasets against published organellar proteomes. For the apicoplast surface, 86 of our candidates (over 68%) overlapped with the dataset from Dong et al. (**Figure 3E**). Their study aimed to identify apicoplast transporters and used a similar approach to ours, fusing TID to APT1, to target the ligase to the organelle surface^31^. To test for spatial specificity, we also compared our surface candidates against the *Plasmodium falciparum* whole-apicoplast proteome generated using a lumenally targeted BirA^57^, after mapping the corresponding *T. gondii* homologs (**Supplemental Table 2**). Reassuringly, only 2% of our apicoplast surface proteome overlapped with this lumenal dataset (**Figure 3E**), confirming high compartment selectivity. We observed a similar pattern across other organellar interfaces: our mitochondrial surface proteome shared only 10% overlap with the mitochondrial matrix proteome reported by Seidi et al.^58^, and our ER surface proteome showed no overlap with candidates generated using the ER lumenal marker TgPDIA3-TurboID^59^. Furthermore, the proportion of integral membrane proteins with one or more transmembrane domains (TMDs) differed by dataset; the ER was the most enriched, with over 75% of candidates predicted to contain at least one TMD (**Figure 3F**). Subsequently, we compared the predicted localization of our candidates with the hyperplexed Localisation of Organelle Proteins by Isotope Tagging (hyperLOPIT) analysis performed in *T. gondii*^46^ (**Figure 3G**). Although the top localization for our apicoplast candidates was the ER (with 27.5% of our proteins, and 6.7% assigned to the apicoplast), this could be explained by the fact that most apicoplast proteins traffic through the ER^60^. As for our other two datasets, 42.5% of our mitochondrial candidates were assigned to the mitochondrial membranes and 63.3% of our ER candidates were assigned to this organelle. Together, these strong positive overlaps, distinct transmembrane enrichment profiles, and minimal overlaps with internal compartments datasets demonstrates that our proximity labeling approach robustly and selectively captures the cytosolic surface of these organelles.

### Identification of a candidate with partial co-localization to both the apicoplast and mitochondrion

Having established our surface proteomes, we sought to identify potential MCS candidates. We reasoned that proteins localized to at least two of our three datasets could represent putative contact-site proteins. Simultaneously, we expanded our organellar surface proteome candidates using a less stringent filtering approach. Specifically, we included proteins that met the following criteria: (i) they were not detected in any of the experiments in the absence of biotin, (ii) at least 5 peptides were detected in at least two of the three biological replicates upon biotin treatment, and (iii) 5 or fewer peptides were detected in the CT-HA^TID^ control strain upon biotin treatment. Using this approach, we expanded our list of candidates to 126 for the apicoplast surface, 63 for the mitochondrial surface, and 168 for the ER (**Figure 4A, Supplemental Table 3**). Using our expanded surface proteomes, we found 40 proteins shared by our apicoplast and ER sets, 12 between our ER and mitochondrial sets and 15 between our mitochondrion and ER sets (**Figure 4B, Supplemental Table 4**). Amongst our apicoplast-ER MCS candidates we found SERCA, known to localize to ER-mitochondrial MCSs in mammalian cells^61^. The list of apicoplast-mitochondrion MCS candidates contains DrpA, which predominantly localizes to the apicoplast and has also been observed colocalizing with the mitochondrion^62^. Similarly, we found LMF1 in the ER and mitochondrial datasets, which is part of a MCS tethering the mitochondrion to the IMC^45^. Among our expanded list of ER-mitochondrial MCS candidates is TgIF2K-A, an ER-resident kinase. Interestingly, TgIF2K-A shares functional features with mammalian PERK^63^. Beyond its role in mediating signal transduction during ER stress, PERK is also an ER-resident component of the lipid trafficking machinery at ER-mitochondria contact sites^64^. Finally, we also found two Rab11 proteins, Rab11A and B, in our three datasets. The very dynamic localization of Rab proteins could explain their presence in our three datasets. Moreover, Rabs are essential for the formation of MCS between the ER and late endosomes in mammals^65^. Altogether, the presence of proteins known to be involved in MCSs gives us high confidence in our approach and provides a robust candidate list for future contact-site investigations. To prioritize candidates for functional characterization, we first filtered our apicoplast-ER MCS candidates by excluding proteins conserved in *Cryptosporidium* species, which lack an apicoplast^66^. We then focused on uncharacterized proteins and further prioritized those with negative phenotype scores^67^, as essential genes are more likely to encode structural or critical MCS components. We also considered one apicomplexan-specific candidate, MtER1. Applying these criteria yielded a list of high-priority MCS candidates **(Figure 4C**). Of note, among those candidates is ApMT2, independently reported to facilitate inter-organelle lipid transport at ER–IMC MCSs^68^, providing strong independent support for our prediction of its role as a MCS candidate. To systematically examine our selected candidates, we use a high-throughput tagging (HiT) strategy^69^ (**Supplemental Figure 4)**. This approach allowed us to endogenously label all our candidates at their C terminus with a 3xTy1epitope tag. Then, we performed IFAs to assess their localizations. Several candidates localized to organelles where we predicted they would be enriched. For instance, MtER2 was confirmed to localize to the mitochondrion through co-localization with TOM40 (**Figure 4D**). ApER1, ApER2, and ApER3 all colocalized with SERCA (**Figure 4E**) verifying their localization to the ER. Interestingly, we also observed several candidates (MtER1, ApMt1, ApMt2, and ApMt3) exhibiting distinct punctate localization patterns, mirroring the characteristic distribution of canonical MCS-resident proteins^1^ (**Figure 4F**). In particular, ApMt1 forms a punctate loop reminiscent of the parasite’s lasso-shaped mitochondrion^70^ showing distinct accumulation at the apical end of the organelle. Based on this unique pattern, we assessed the co-localization of ApMt1 with the apicoplast (**Figure 4G**) and the mitochondrion (**Figure 4H**). The partial co-localization observed with both organelles strongly suggested a potential role in organelle contact sites, prompting us to select this candidate for further characterization.

**Figure 4:**
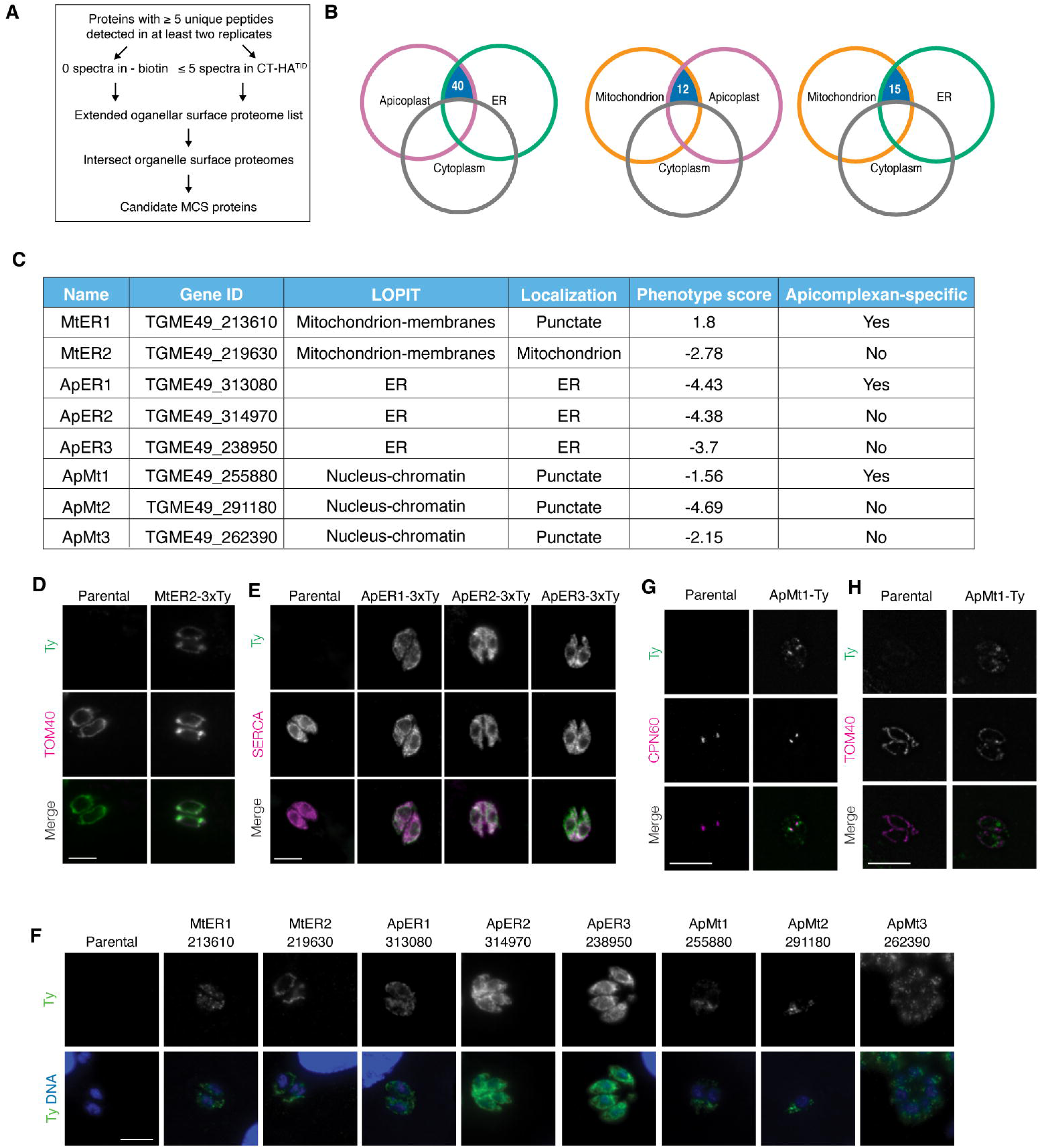
Localization of MCS candidates. **(A)** Experimental design used to expand the initial surface proteome candidate list. **(B)** Venn diagrams showing the number of shared candidate proteins among the apicoplast, mitochondrial, and ER surface datasets. **(C)** Table summarizing the selected MCS protein candidates identified in this study. **(D)** Intracellular parasites expressing MtER2-3xTy were fixed and stained for Ty (green) and the mitochondrial marker TOM40 (magenta). **(E)** Intracellular parasites expressing ApER1-3xTy, ApER2-3xTy, or ApER3-3xTy were fixed and stained for Ty (green) and the ER marker SERCA (magenta) **(F)** Intracellular parasites expressing epitope-tagged MCS candidate proteins were fixed and stained for Ty (green) and DAPI (blue). **(G–H)** Parasites expressing ApMt1-Ty were fixed and stained for Ty (green) along with either the apicoplast marker CPN60 **(G)** or the mitochondrial marker TOM40 (magenta) **(H)**. Scale bars: 5 µm.

### ApMt1 depletion impairs parasite viability and disrupts putative apicoplast-mitochondrion contact sites

To further characterize ApMt1’s potential role in the apicoplast-mitochondrion MCSs, we generated an Ty-tagged, conditional knock-down strain using the DiCre mediated U1 gene silencing approach^71^ as this gene is predicted to be essential^67^ (**Supplementary figure 5A**). Western blot analysis confirmed ApMt1 depletion 72 hours following a 2-hour rapamycin treatment (**Figure 5A**). Correct localization of the protein and subsequent loss of Ty signal upon rapamycin treatment was also observed by IFA (**Supplementary figure 5B**). Plaque assays confirmed that loss of ApMt1 resulted in a significant decrease in plaque area, indicating that silencing of the protein has a negative impact on parasite viability (**Figure 5B–C**). Finally, to assess whether ApMt1 plays a role in mediating MCSs between the apicoplast and mitochondrion, we performed super-resolution imaging followed by quantitative spatial analysis. Although we observed no gross defects in apicoplast or mitochondrial morphology (corroborated by similar organellar surface area and volume measurements (**Supplementary figure 5C-F**)) ablation of ApMt1 led to a significant decrease in signal co-localization between the two organelles when compared to the parental strain (**Figure 5D–E**). This spatial separation indicates that the apicoplast and mitochondrion are significantly farther apart in the absence of ApMt1, confirming that ApMt1 is a functional tether at the apicoplast–mitochondrion MCS. Although we did not find a significant difference between DMSO and rapamycin treated ApMt1^cKD^, this is likely due to the well-characterized leakage of the conditional knockdown system^71^ and may indicate that expression level of ApMt1 is important for apicoplast-mitochondrion contact spacing. Collectively, our results support the existence of mitochondrion–apicoplast MCSs and demonstrate that ApMt1 is required to maintain their proper contact.

**Figure 5:**
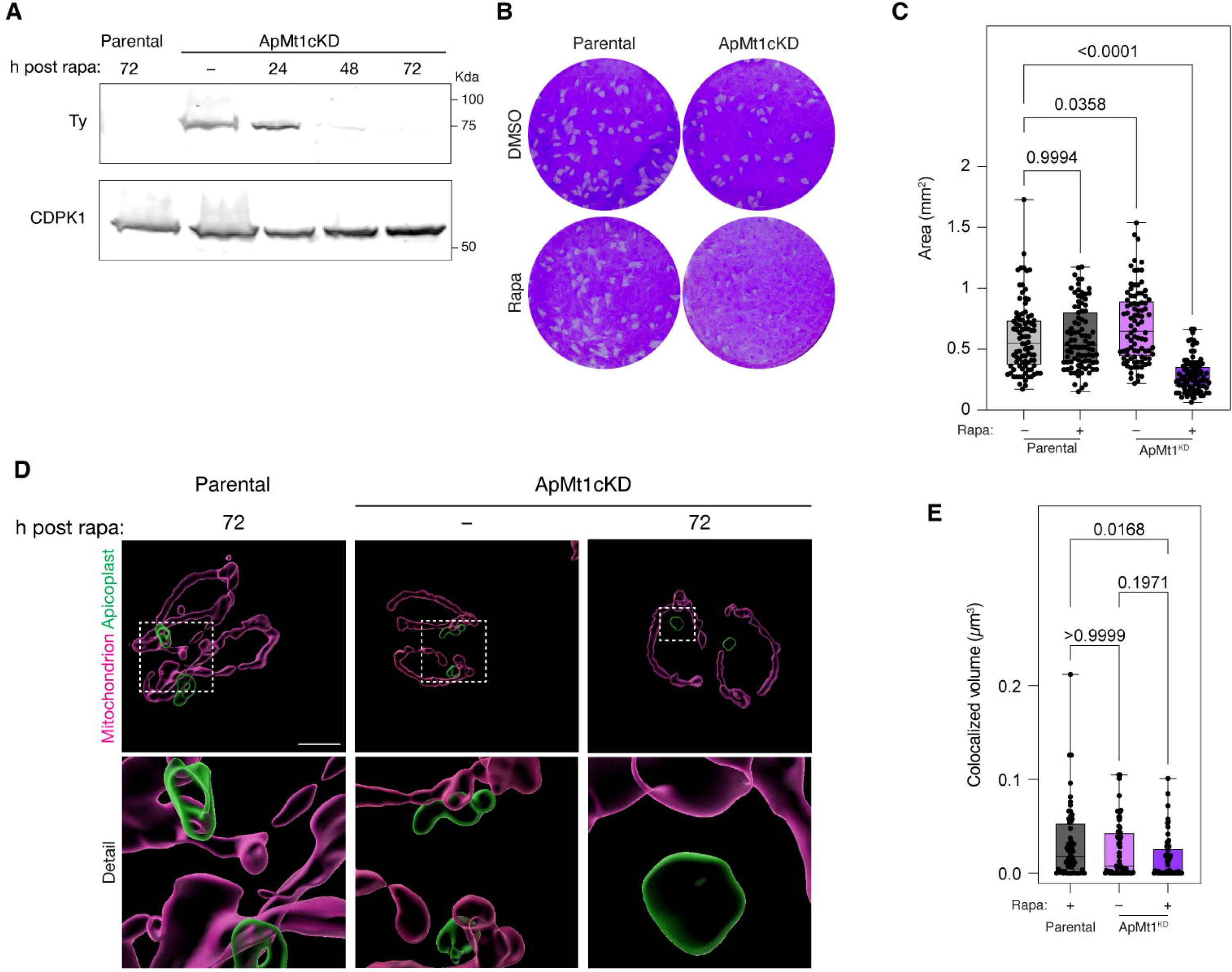
Loss of ApMt1 reduces parasite viability and apicoplast-mitochondrion MCSs. **(A)** Parental and ApMt1^cKD^ strains were pulsed with DMSO or rapamycin at the indicated time points before collection of lysates. Samples were separated via SDS-PAGE and the resulting blot was probed with antibodies for Ty (upper panel) and CDPK1 as a loading control (lower panel). Data are representative of three biological replicates. **(B)** Parental or ApMt1^cKD^ parasites were allowed to invade confluent 6-well plates for two hours with DMSO or Rapamycin pulse before washing. Plates were incubated for 8 days prior to fixing and staining with crystal violet. Images are representative of 3 biological replicates. **(C)** Plaque area was measured for 90 plaques across 3 biological replicates in FIJI. **(D)** Parental or ApMt1^cKD^ parasites were pulsed with DMSO or rapamycin 72 hours before fixation and subsequent staining with the mitochondrial TOM40 (magenta) and the apicoplast marker CPN60 (green) for super-resolution imaging and 3D model reconstruction in IMARIS software. Representative images of 3 biological replicates. Scale bars are 2 µm. **(E)** IMARIS software was used to calculate the overlapped volume of the two organelles. A total of 59 individual parasites per condition were measured across 3 biological replicates. Imaging and analysis were performed blinded.

## Discussion

Membrane contact sites (MCSs) represent crucial hubs of organelle communication defined by specific proteomic compositions^1^. Mapping the proteomes of these domains is notoriously difficult because standard isolation protocols generally disrupt the interactions that hold apposing organellar membranes together. Here, we employed proximity biotinylation to overcome these technical limitations. A key advantage of this approach is that it enables *de novo* candidate discovery without requiring prior knowledge of MCS components—a particularly powerful capability given our rudimentary understanding of inter-organelle contact sites in *T. gondii* and other apicomplexans^24^.

After validating the membrane localization and topology of our organelle TID handles, we performed proteomic analyses against a cytosolic spatial reference to establish stringent surface proteomes for the apicoplast, mitochondrion, and ER. Our results align with previously published localizations and uncover novel predictions that highlight several proteins as key MCS candidates, including TgVPS13A, (which we designated ApMT2), a candidate whose role at ER–IMC contact sites was independently confirmed in a study published while our manuscript was in preparation^68^. VPS13-family proteins function as bulk lipid transporters that bridge organellar membranes through interactions with organelle-bound adaptors^72^, with adaptor identity dictating organellar specificity^73^. Although ApMT2/TgVPS13A was significantly enriched in our ER biotin-treated samples over unlabeled controls, it did not meet our high-stringency threshold when filtered against the cytosolic reference. Nevertheless, its discovery in our apicoplast and mitochondrial surface proteomes, combined with the flexible nature of VPS13 adaptors, suggests that ApMT2/TgVPS13A may play a broader role in lipid transport across multiple contact sites, including those between the mitochondrion and apicoplast.

While many of our candidates matched their reported subcellular locations^46^, others displayed a punctate pattern despite being assigned to the nucleus by hyperLOPIT. Because hyperLOPIT measures steady-state protein distributions, proteins that shuttle between or localize to multiple organelles cannot always be unambiguously classified^74^. Indeed, many MCS proteins form discrete punctate, dot-like patterns due to tight clustering at narrow zones where organelles interact^1^. Given the dynamic nature of these contact sites, it is plausible that hyperLOPIT assignments may mask or misclassify proteins with transient or dual-organelle distributions.

One of these candidates, ApMT1, exhibited a punctate pattern reminiscent of the parasite’s lasso-shaped mitochondrion^70^ but with a distinct accumulation near the apical pole of the nucleus, a zone where the apicoplast resides. ApMT1 partially co-localized with both organelles, and conditional depletion revealed that it is essential for parasite survival. Crucially, super-resolution microscopy demonstrated that ApMT1 knockdown caused a significant reduction in signal co-localization between the apicoplast and mitochondrion. This reduction indicates that the two organelles shift apart in the absence of ApMT1, establishing ApMT1 as a functional tether at apicoplast–mitochondrion contact sites. Although physical proximity between the mitochondria and the apicoplast was reported six decades ago^75^, its molecular identity and precise function remained unknown. However, this interaction is likely mediated by MCSs to facilitate metabolic exchange, its molecular identity and precise function remained unknown, although it likely facilitates metabolic exchange and mediated by MCSs. Our findings establish ApMT1 as a component mediating this inter-organellar contact. Secondary structure homology analysis using HHpred^76^ indicates that the N-terminal region of ApMt1 shares structural homology with the membrane-interacting motifs of the pore-forming toxins ostrolysin and fragaceatoxin C^77,78^, suggesting that this domain may directly insert into organellar membranes. While future biochemical and structural characterization will be necessary to test this model and define its precise mechanism of membrane integration, ApMT1 represents a major step forward in defining the molecular architecture of apicomplexan contact sites and, to our knowledge, the first apicoplast-mitochondrion MCS candidate.

Taken together, our findings demonstrate the power of proximity labeling for identifying MCS components in *Toxoplasma gondii*. In particular, pairing proximity labeling with a cytosolic spatial reference provides a robust, unbiased framework for generating organellar surface proteomes and uncover novel contact site candidates without prior candidate knowledge. We also demonstrate that organelle-surface handles can be readily adapted for diverse cellular compartments. This strategy provides a framework for studying not only other subcellular compartments but also other apicomplexan species, including *Plasmodium falciparum*, the causative agent of malaria, to systematically characterize MCS composition across this major group of human pathogens.

---

**Supplementary figure 1:**
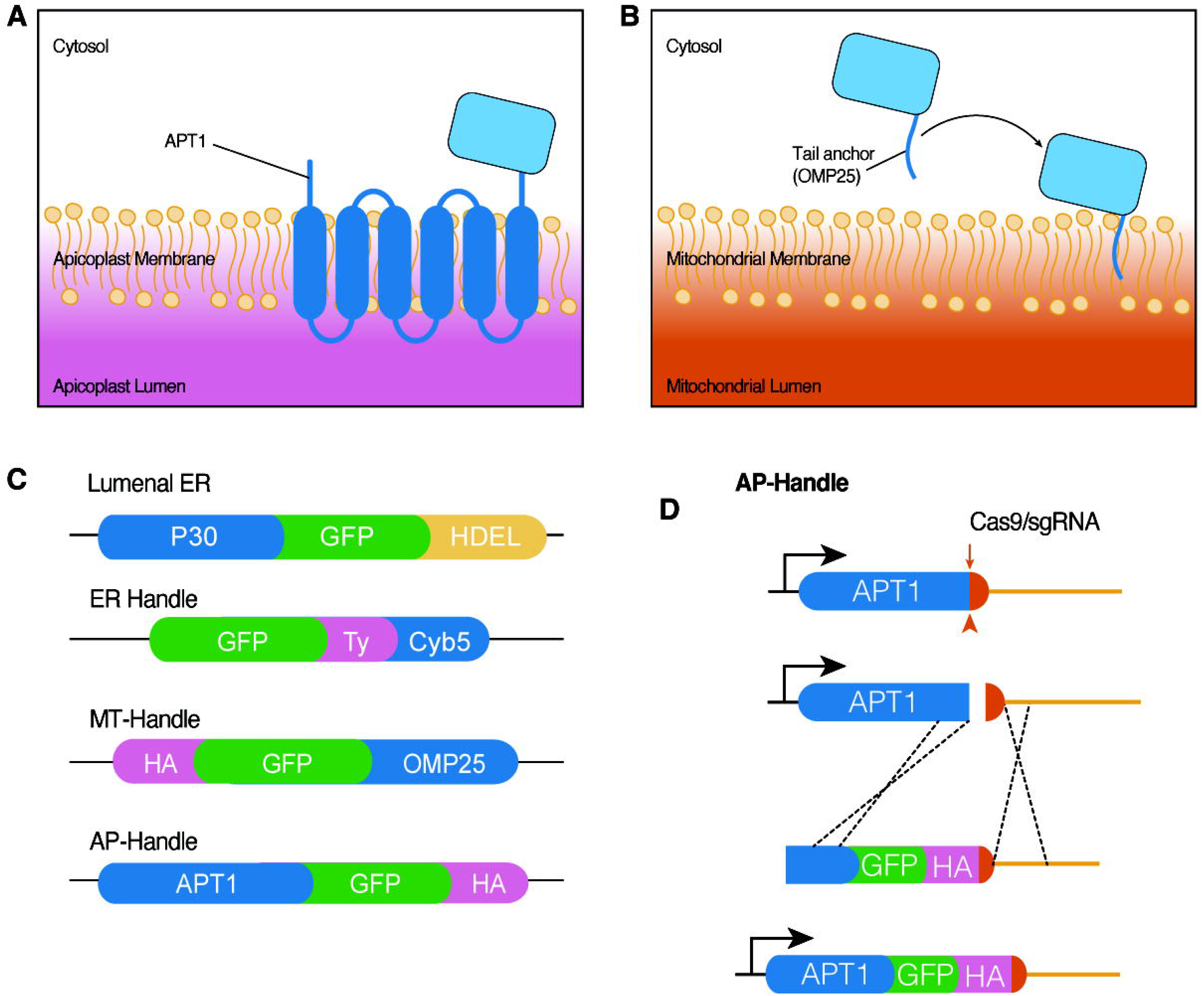
Organelle-surface targeting strategies. **(A)** Targeting proteins to the outer apicoplast membrane via C-terminal tagging of APT1. **(B)** Targeting proteins to the mitochondrial surface using the transmembrane domain of OMP25. (**C**) Diagrams of organelle-surface and ER luminal targeting constructs used in this study. (**D**) Schematic depicting the strategy used to tag APT1 with GFP.

**Supplementary figure 2:**
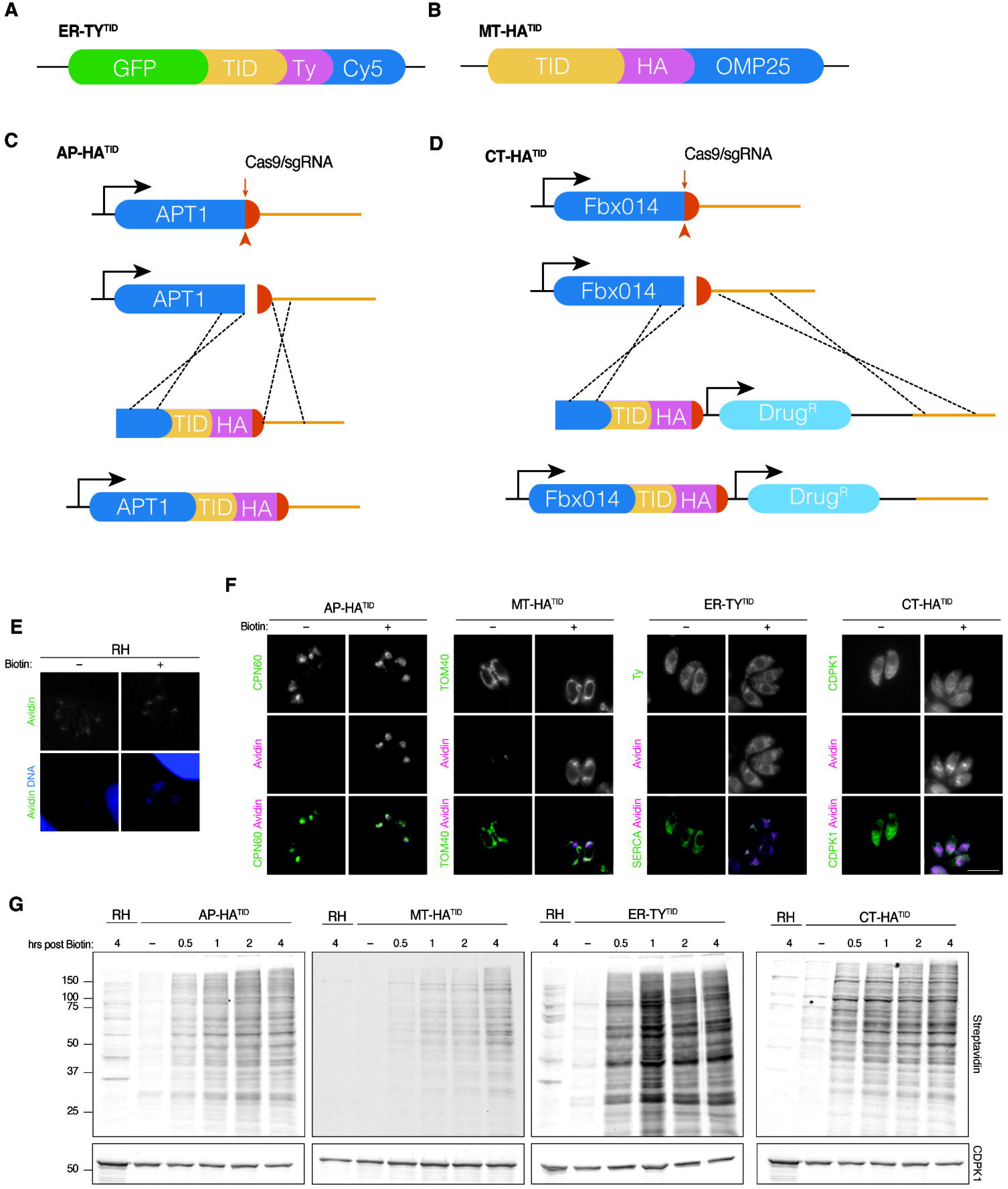
Generation and validation of TurboID expressing lines. **(A-B)** Schematic of the plasmid constructs used to generate the ER-TY^TID^ **(A)** and MT-HA^TID^ **(B)** organellar handles. **(C-D)** Schematic representation of the CRISPR/Cas9-mediated tagging strategy to generate AP-HA^TID^ **(C)** and CT-HA^TID^ **(D)**, respectively. **(E)** RH parasites incubated with 50 µM biotin or DMSO (vehicle control) for 4 hours were fixed and stained with Avidin (green) and DAPI (blue). **(F)** Immunofluorescence assays of AP-HA^TID^, MT-HA^TID^, and CT-HA^TID^ incubated with 50 µM biotin or DMSO for 4 hours prior to fixation and staining with Avidin (green) and the organelle markers CPN60, TOM40, Ty, or CDPK1 (magenta) to label the apicoplast, mitochondrion, and cytoplasm, respectively. Ty signal from the ER-Ty^TID^ strain was used to confirm labelling on the ER. **(G)** Parental and parasite lines expressing the specified TurboID constructs (AP-HA^TID^, MT-HA^TID^, ER-Ty^TID^ and CT-HA^TID^) were treated with 50 μM biotin or DMSO for the indicated amount of time. Whole-cell lysates were separated by SDS-PAGE and blotted with streptavidin to visualize protein biotinylation (upper panel). CDPK1 served as a loading control (lower panel).

**Supplementary figure 3:**
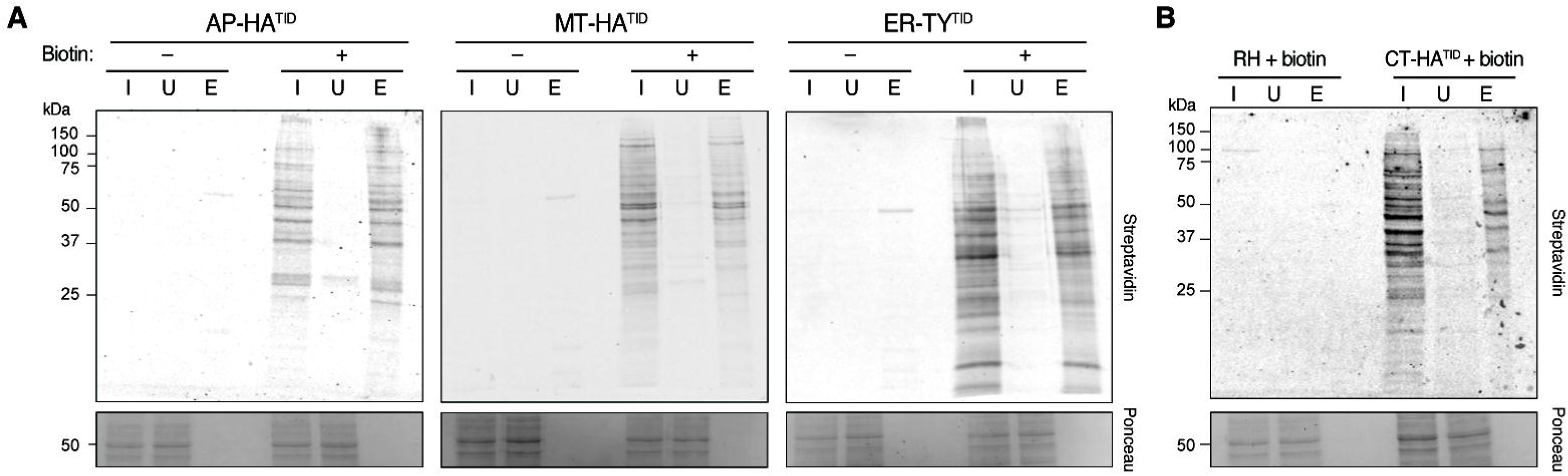
Enrichment of biotinylated proteins. (**A-B**) Parasite lines expressing AP-HA^TID^, MT-HA^TID^, or ER-Ty^TID^ (**A**) along with RH and CT-HA^TID^ parasites (**B**) were incubated with 50 uM biotin or DMSO for 4 hours prior to collection, cell lysis and streptavidin pull-down. The Input (I), Unbound (U), and Eluted (E) fractions were separated via SDS-PAGE. Ponceau stain is included to show equal loading of total protein (lower panel) and the blot was probed with streptavidin to confirm successful enrichment of biotinylated proteins.

**Supplementary figure 4:**
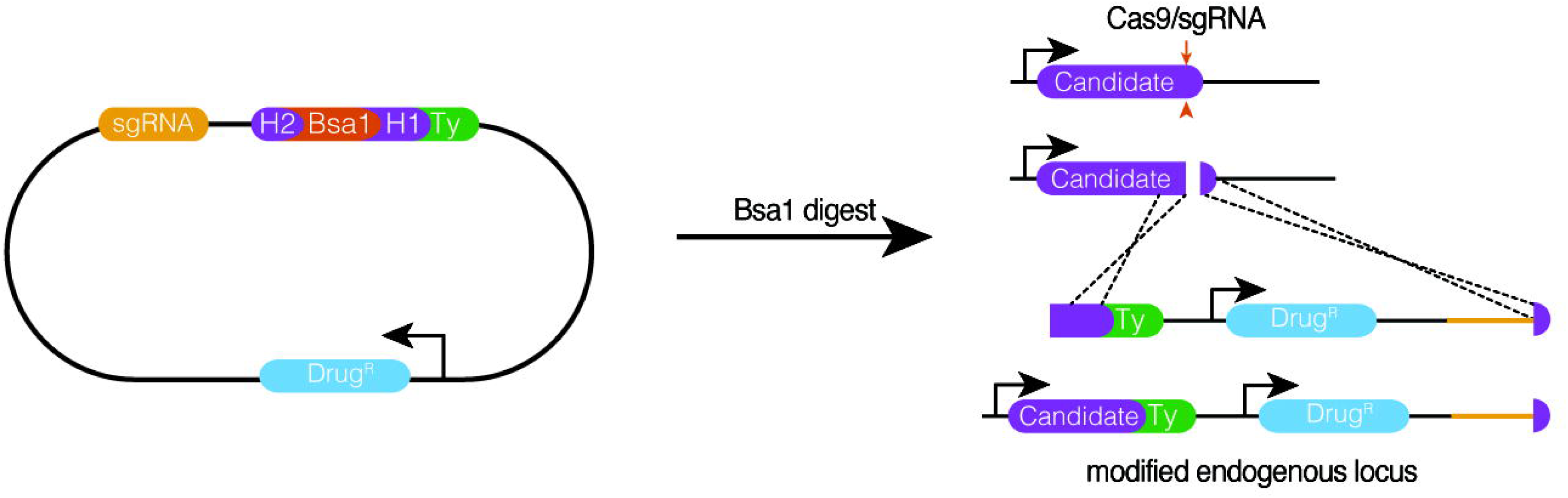
Strategy used to tag MCS candidate proteins. Schematic of the CRISPR/Cas9 mediated C-terminal Hit tagging of candidate proteins.

**Supplementary figure 5:**
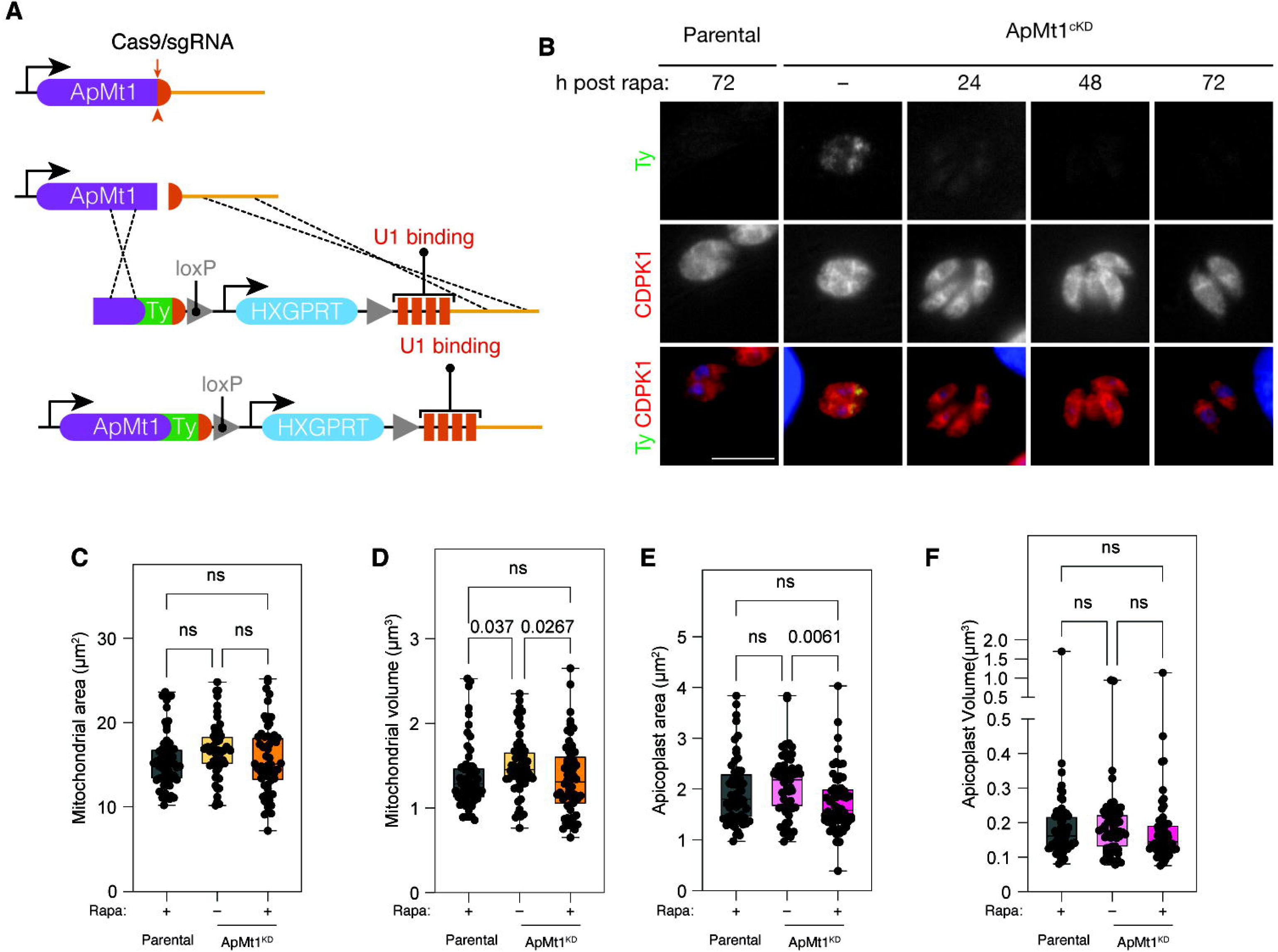
Characterization of ApMt1^cKD^. **(A)** Schematic of the ApMT1 locus and the strategy to generate the ApMt1^cKD^ strain. **(B)** Following treatment with rapa or vehicle (–), intracellular ApMt1^cKD^ parasites were fixed and stained for Ty (green), CDPK1 (red), and DAPI (blue). **(C-F)** Quantification of mitochondrial (**C, D**) and apicoplast (**E, F**) surface area and volume using Imaris. A total of n = 59 parasites across 3 biological replicates were used and the analysis was blinded. Groups were compared using a Krustal-Wallis test followed by Dunn’s test for post-hoc multiple comparison.

## Acknowledgements

This work was supported by the National Institutes of Health (K.V.P.: T32AI060546, D.H.: R35GM150794)

We would like to thank Dr. Muthugapatti K. Kandasamy and the Biomedical Microscopy Core at the University of Georgia, as well as Julie Nelson and Juan Bustamante at the CTEGD Cytometry Shared Resource Laboratory, for their technical expertise and access to imaging and cell-sorting equipment. We are grateful to the Mass Spectrometry & Biopolymers Core at the Koch Institute’s Robert A. Swanson (1969) Biotechnology Center, specifically Richard P. Schiavoni, for invaluable technical support with mass spectrometry experiments. We also thank Boris Striepen, Sebastian Lourido, and Tyler Smith for sharing antibodies and reagents, as well as Silvia Moreno and the members of the Moreno laboratory for providing antibodies, reagents, and indispensable feedback on this work. We acknowledge Brittany Henry and Nichole Khamsa for their assistance in generating the AP-HA^TID^, MT-HA^TID^ and CT-HA^TID^ strains. Last but not least, we thank VEuPathDB and all contributors to this resource.

## Author contributions

Conceptualization, formal analysis, funding acquisition, investigation, methodology and writing: K.V.P. and D.H

## Declaration of interests

The authors declare no competing interests

## Materials and Methods

### Parasite culture

*T. gondii* tachyzoites were maintained in human foreskin fibroblasts (HFFs) (ATCC, cat. no. SCRC-1041). Host cells and parasites were cultured in DMEM (Thermo Fisher Scientific, cat. no. 11965118) with 2mM glutamine (GeminiBio, cat. no. 400-106), 0.1µg/mL Gentamicin (Thermo Fisher Scientific, cat. no. 15710072), and 3% heat-inactivated fetal calf serum (IFS, Gemini Bio, cat. no. B7002S) at 37°C with 5% CO_2_.

### Plasmid and strain generation

Oligonucleotides were synthesized by Integrated DNA Technologies (IDT). PCRs and assembly steps were performed using Q5 High-Fidelity 2× Master Mix and NEBuilder HiFi DNA Assembly Master Mix (New England Biolabs). All primers and plasmids are listed in **Supplementary Table 5**. To generate the apicoplast surface handle, the endogenous locus of APT1 (TGME49_261070) was modified. An sgRNA targeting the C terminus of APT1 (P1 and P2) was inserted into *BsaI*-digested pU6-Universal plasmid^79^ using Gibson assembly. A repair template consisting of GFP-3xHA with homology regions to the 3’UTR of APT1 was amplified from the OMP25-GFP plasmid^80^ (named pOMP25-GFP) using P3 and P4. Approximately 15 μg of this repair template and 50 μg of the pU6-Universal plasmid encoding Cas9 containing the C-terminal targeting sgRNA were transfected into RH/Δku80/Δhx parasites as described previously. Four days after transfection, freshly lysed GFP-positive parasites were sorted and subcloned into a 96 well plate using a Cytek Aurora CS (Cytek Biosciences, Bethesda, MD). Successful APT1 tagging was confirmed via live microscopy and immunofluorescence. To generate the ER surface handle (pER-GFP), a plasmid containing the eGFP coding sequence followed by an in-frame single Ty epitope and the last 105 bp of the 3’ end of the Cytochrome b5 (TGME49_276110) was generated. To do so, the backbone (p-Tub1) was digested with Nhe1 and Mfe1, and the eGFP fragment was PCR amplified from our GFP-containing plasmid p-GFP using primers P5 and P6. The Ty-Cyb5 fragment was PCR amplified from parasite genomic DNA using primers P7 and P8 and the resulting linearized vector and fragments were assembled via Gibson assembly. GFP-ER-expressing parasites were generated by transfecting 100 µg of pER-GFP into RH parasites as previously described^79^ and selecting with 40 µM chloramphenicol. After selection, the population was subcloned by serial dilution to obtain monoclonal populations and validated using live imaging, IFA and Western blotting. The luminal GFP-ER plasmid was generated by amplifying the 5’UTR of DHFR from the DHFR-SAG4-TetO7-3xTy plasmid (a kind gift from the Moreno Lab) with primers P9 and P10, while the P30 domain was amplified from parasite genomic DNA with primers P11 and P12, eGFP-HDEL was amplified from p-GFP using P13 and P14, and pTub1 was digested with Nru1 and Mfe1 prior to Gibson assembly, generating p-eGFP/HDEL. To allow for simultaneous expression of the mCherry-GFPnanobody in the same construct, p-eGFP/HDEL was then digested with Kpn1 and Age1 and Gibson assembled with a fragment containing the mCherry-GFPnanobody PCR-amplified from the GFP-nanobody plasmid^80^ using primers P15 and P16 to generate the pGFP-HDEL/GFP-Nb plasmid.

A similar approach was used to the TurboID-expressing handles, which are derivatives of our GFP-expressing constructs. AP-HA^TID^ was generated by targeting the C terminus of APT1 with the same pU6-Universal plasmid containing the sgRNA targeting the C terminus of the gene (P1 and P2). A repair template consisting of TurboID-3xHA with homology regions to the 3’UTR of APT1 was amplified from the 3HA-TurboID pLIC plasmid (a kind gift from the Moreno lab) using P17 and P18. Approximately 20 μg of this repair template and 50 μg of the pU6-Universal plasmid encoding Cas9 containing the C-terminal targeting sgRNA were transfected into RH/Δku80/Δhx parasites as described previously (Sidik et al., 2014). Successful integration of the repair template was assessed via P19 and P20, and the population was subcloned via serial dilution. Clonal lines were screened for the presence of TID via PCR using P17 and P18. Presence of HA-tagged TID was confirmed via immunofluorescence. The mitochondrial MT-HA^TID^ plasmid was generated by modifying pOMP25-GFP. The backbone was linearized using NheI and AvrII. Two PCR fragments, TID (amplified from the 3HA-TurboID pLIC plasmid using primers P21 and P22) and the OMP25 localization signal (amplified from pOMP25-GFP using primers P23 and P24), were then inserted into the digested vector backbone via Gibson assembly. The ER-Ty^TID^ plasmid was generated by first amplifying 4 different fragments: the 5’UTR of MIC2 from genomic DNA using P25 and P26; eGFP from pER-GFP plasmid using P27 and P28; TurboID from the MT-HA^TID^ plasmid using P29 and P30 and Ty-Cy5 from PER-GFP/pKP001 using P31 and P32. The fragments were inserted into pTub1 linearized with Nru1 and MfeI by Gibson assembly. To generate the ER-Ty^TID^ strain, 100 µg of pER-Ty^TID^ was transfected into RH strain parasites as previously described^79^. Two days after transfection, the parasites were sorted using a Cytek Aurora CS (Cytek Biosciences, Bethesda, MD) to enrich for GFP positive parasites before selecting with 40 µM chloramphenicol. After selection, the population was subcloned by serial dilution to obtain monoclonal populations and validated using IFA and Western blotting. The CT-HA^TID^ plasmid was generated using the HiT strategy^69^. The V5-T2A-mKate2 Hit vector (a kind gift from the Lourido lab) was digested with AvrII and PacI, and a payload (gBlock), P33, containing the sgRNA targeting FBXO14 (TGME49_259880), three in-frame HA epitope tags, and TID was inserted into the digested V5-T2A-mKate2 by Gibson assembly. The resulting plasmid was then linearized with BsaI and cotransfected with the Cas9-expression plasmid pSS014 as previously described^69^. Successfully modified parasites were selected using 3 µM pyrimethamine prior to subcloning via serial dilution to obtain a monoclonal population. Clones were validated by IFA and Western blotting.

The HiT strategy was also used to tag the selected MCS candidates. First, a HiT vector with a 3x-Ty1 tag payload (pHiT-3xTy) was generated by cloning a gBlock containing a 3xTy1 tag (P34) into the V5-T2A-mKate2 HiT vector by Gibson assembly. For each gene, a payload consisting of the sgRNA targeting the respective 3’ end of each candidate and a 40 bp homology region immediately upstream and downstream of the stop codon (P36-51) was cloned into the pHiT-3xTy vector by Gibson assembly. RH/Δku80/Δhx parasites were transfected with 50 µg of Bsa1 digested vector and 50 µg of the Cas9 plasmid pSS014^69^ prior to selection using 3 µM pyrimethamine. The resulting mixed populations were analyzed by immunofluorescence assay to determine the subcellular localization of the MCS candidates.

To generate the ApMt1^cKD^ strain, a sgRNA targeting the C-terminus of ApMt1 (P51 and P52) was Gibson assembled into the Bsa1 digested pU6-Universal plasmid (Addgene, cat. no. 52694). A repair template containing 3xTy1-loxP-HXGPRT cassette-loxP-U1 flanked by 40 bp of homology immediately upstream and downstream of the stop codon was amplified from the 3xTy-U1 vector using P53 and P54. RH DiCre_T2A Δku80Δhxgprt^81^ (parental) tachyzoites were then co-transfected with ∼10 µg of the repair template and ∼30µg of the pU6-Universal plasmid containing the sgRNA and selected with 50mg/mL xanthine and 25 mg/mL mycophenolic acid. The resulting mixed population was subcloned via serial dilution to obtain a monoclonal population, which was validated by PCR (P55 and P56), IFA, and Western blotting.

### Immunofluorescence assays

HFFs were seeded on glass coverslips and infected with 20-50 µL of extracellular tachyzoites once confluent. While still intracellular, coverslips were fixed with 4% paraformaldehyde in PBS for 15 minutes at 4°C, permeabilized with 0.25% Triton-X in PBS for 8 minutes at RT, and blocked in 5% IFS and 5% normal goat serum (NGS) diluted in PBS. Primary antibody staining was conducted in blocking solution for 1-4 hours using mouse anti-calumenin (a kind gift from the Moreno lab); mouse anti-Ty1 ^82^; rabbit anti-HA (Abcam, cat. no. ab9110); guinea pig anti-SERCA^83^; guinea pig anti-CDPK1 (a kind gift from the Lourido lab), rabbit anti-CPN60 or rabbit anti-TOM40 (both kind gifts from the Striepen lab) as indicated. Secondary antibody staining was diluted in blocking solution (5% IFS, 5% NGS in PBS) and incubated for 1 hour at RT. The following antibodies were used: Alexa Fluor 488-conjugated goat-anti-mouse (Invitrogen, cat. no. A32723); Alexa Fluor 488-conjugated goat-anti-rabbit (Invitrogen cat. no. A32731); Alexa Fluor 488-conjugated goat-anti-guinea pig (Invitrogen, cat. no. A11073); Alexa Fluor 594-conjugated goat-anti-guinea pig (Invitrogen, cat. no. A11076); Alexa Fluor 647-conjugated goat-anti-mouse (Invitrogen, cat. no. A32728); Alexa Fluor 647-conjugated goat-anti-rabbit (Invitrogen, cat. no. A32733); or Alexa Fluor 647-conjugated goat-anti-guinea pig (Invitrogen, cat. no. A21450). Hoescht (Santa Cruz Biotechnology, cat. no. sc-394039) was used to label nuclei and was diluted in blocking solution alongside the secondary antibodies. Coverslips were mounted on slides using ProLong diamond (Thermo Fisher Scientific, cat. no. P36961), and images were acquired on an ECHO Revolve microscope and the ECHO Pro application. For co-localization of ApMt1, images were acquired on a Delta Vision microscope (Olympus, IX-71) and were deconvolved using the softWoRx deconvolution software. All image analysis and processing were conducted using Fiji, Adobe Photoshop 2024-2026, and Adobe Illustrator 2024-2026.

### Live Imaging

To determine the topology of the organelle handles, parasite strains individually expressing each candidate handle, along with a parental control strain, were transfected with 100 µg of pGFP-nanobody plasmid. As a control for ER luminal GFP localization, RH parasites were transfected with pGFP-HDEL/GFP-Nb plasmid. Immediately following transfection, 20-50 µL of tachyzoites were inoculated onto human foreskin fibroblast (HFF) monolayers in 35mm glass-bottom dishes (Mattek) and allowed to recover for 24 hours prior to live-cell imaging. Images were acquired using an ECHO Revolve microscope with the ECHO Pro application. Image processing and analysis were performed using Fiji, Adobe Photoshop 2024, and Adobe Illustrator 2024.

### Western Blotting

To prepare samples for Western blot analysis, parasite pellets (approximately 5×10^7^ parasites) were resuspended in 2X Laemmli sample buffer (20% glycerol, 5% 2-mercaptoethanol, 4% SDS, 0.02% bromophenol blue, 120 mM Tris-HCl, pH 6.8) and boiled at 100°C for 5 min. Precision Plus Protein Dual Color Standard (Bio-Rad, Cat. No. 1610374) was used as a molecular weight marker. Proteins were resolved by SDS-PAGE and transferred onto nitrocellulose membranes. Membranes were probed with mouse anti-Ty and guinea pig anti-CDPK1 primary antibodies. Secondary detection was performed using donkey anti-mouse IgG conjugated to IRDye 680RD (LICORBio, Cat. No. 926-68070) and goat anti-guinea pig IgG conjugated to IRDye 800CW (LICORBio, Cat. No. 926-32411). Blots were imaged using an Odyssey infrared imaging system (LICORbio).

### Enzymatic activity validation of TurboID

For IFA validation, coverslips seeded with HFFs and infected with AP-HA^TID^, MT-HA^TID^, ER-Ty^TID^, or CT-HA^TID^ tachyzoites were incubated with 50 µM biotin or DMSO (vehicle control) for 4 hours at 37°C prior to fixation and processing as described previously. Primary antibodies, namely rabbit anti-CPN60, rabbit anti-TOM40, mouse anti-Ty1, and guinea pig anti-CDPK1, were used as indicated in the figure captions, followed by secondary detection using Alexa Fluor 488-conjugated goat anti-mouse (Invitrogen, Cat. No. A32723), Alexa Fluor 488-conjugated goat anti-rabbit (Invitrogen, Cat. No. A32731), Alexa Fluor 488-conjugated goat anti-guinea pig (Invitrogen, Cat. No. A11073), and NeutrAvidin Texas Red conjugate (Thermo Fisher Scientific, Cat. No. A2665). For Western blot validation, parasites were incubated with 50 µM biotin or DMSO for the indicated times prior to lysis and Western blotting as previously described. Blots were probed with guinea pig anti-CDPK1 followed by goat anti-guinea pig IgG conjugated to IRDye 680RD (LICORBio, Cat. No. 926-68077) and IRDye 800CW Streptavidin (LICORBio, Cat. No. 926-32230). Proteins were visualized using an Odyssey infrared imager (LICORBios).

### Enrichment of biotinylated proteins

RH, AP-HA^TID^, MT-HA^TID^, ER-Ty^TID^, and CT-HA^TID^ tachyzoites were used to infect T75 flasks containing confluent hTERT-immortalized cells (human telomerase reverse transcriptase, a gift from the Moreno lab) and allowed to grow for 36 hours. For the AP-HA^TID^, MT-HA^TID^ and ER-Ty^TID^ parasites, 10 flasks of each strain were incubated with 50 µM biotin (5 flasks) or DMSO (5 flasks) for 4 hours. The RH and CT-HA^TID^ strains were incubated with 50 µM biotin (5 flasks for each strain) for 4 hours. Biotinylation was stopped by placing flasks on ice and washing 3x with ice-cold PBS. The remaining intracellular parasites were mechanically lysed by passing through a 30-gauge needle and filtered to remove host debris. Streptavidin pulldown was performed as previously described^84^. In brief, 1×10^8^ parasites were lysed in 500 mL cold RIPA buffer (Thermo Fisher Scientific, cat. no. PI89901) supplemented with 1× HALT Protease and Phosphatase Inhibitor (Thermo Fisher Scientific, cat. no. PI78440) for 10 mins on ice with occasional vortexing and clarified by centrifugation in 4°C tabletop centrifuge at 12000 rpm. The resulting lysates were added to 100 mL of streptavidin beads and incubated at 800 rpm at 4°C for 1 hour. The beads were then washed and sent for on-bead digestion.

### Mass spectrometry

Proteins bound to streptavidin beads were reduced with 10mM dithiothreitol for 1h at 56°C and then alkylated with 20mM iodoacetamide for 1h at 25°C in the dark. Then proteins were then digested with Modified Trypsin in 100mM ammonium bicarbonate, pH 8 at 25°C overnight. Trypsin activity was halted by addition of formic acid (99.9%) to a final concentration of 5%. Peptides were desalted using Pierce Peptide Desalting Spin columns then vacuum centrifuged.

The tryptic peptides were separated by reverse phase HPLC using a Thermo PepMap RSLC C18 column (2µm tip, 75µmx50cm) over a 90-minute gradient before nano electrospray using a Orbitrap Exploris 480 mass spectrometer (Thermo). Solvent A was 0.1% formic acid in water and solvent B was 0.1% formic acid in acetonitrile. The gradient conditions were 1% B (0-10 min at 300nL/min) 1% B (10-15 min, 300 nL/min to 200 nL/min) 1-7% B (15-20 min, 200nL/min), 7-25% B (20-54.8 min, 200nL/min), 25-36 B (54.8-65 min, 200nL/min), 36-80% B (65-65.5 min, 200 nL/min), 80% B (65.5-70 min, 200nL/min), 80-1% B (70-70.1 min, 200nL/min), 1% B (70.1-90 min, 200nL/min).

The Thermo Orbitrap Exploris 480 mass spectrometer was operated in a data-dependent mode. The parameters for the full scan MS were: resolution of 120,000 across 375-1600 m/z and maximum IT 25 ms. The full MS scan was followed by MS/MS for as many precursor ions in a two second cycle with a NCE of 28, dynamic exclusion of 20s and resolution of 30,000.

Raw mass spectral data files (.raw) were searched using Sequest HT in Proteome Discoverer (Thermo) the *Toxoplasma gondii* ME49 protein database (Uniprot) and a made in house database of common contaminants (e.g., keratin, trypsin). with the following search parameters: 10ppm mass tolerance for precursor ions; 0.02 Da for fragment ion mass tolerance; 2 missed cleavages of trypsin; fixed modification were carbamidomethylation of cysteine, variable modifications were methionine oxidation, methionine loss at the N-terminus of the protein, acetylation of the N-terminus of the protein and also Met-loss plus acetylation of the protein N-terminus. Mascot search results were imported into Scaffold (Proteome Software), applying a minimum peptide threshold of 95% confidence prior to downstream surface proteomone analysis using custom R scripts.

### Super-resolution imaging

Intracellular RH DiCre_T2A Δku80Δhxgprt (parental) or ApMt1^cKD^ parasites were incubated with 50nM rapamycin or DMSO (vehicle control) for 2 hours, washed twice with PBS, and replenished with fresh culture medium. After 72 hours, parasites were fixed and processed for immunofluorescence assay (IFA) as described previously. For the primary antibody staining, Rabbit anti-CPN60 and guinea pig anti-TOM40 were used, followed by secondary staining with Alexa Fluor 488-conjugated goat-anti-rabbit (Invitrogen cat. no. A32731), Alexa Fluor 594-conjugated goat-anti-guinea pig (Invitrogen, cat. no. A11076) and Hoechst to label cell nuclei. Images were acquired with Ziess ELYRA S1 system with an EM-CCD camera (Andor iXon) and a 100x/ oil immersion objective using the three-phase setting. A total of 46 z-stacks were taken in 0.11 µm increments for a total of 5.06 µm. Structured illumination was performed using the accompanying ZEN 2011 software with SIM analysis module, and 3D reconstructions were assembled in the IMARIS 10.2 software for area and volumetric measurement of the organelles to assess morphology. Volumes of overlapped surfaces were measured in the IMARIS 10.2 software to assess co-localization, corresponding to MCSs. Slides were blinded prior to imaging and analysis. Significance was calculated using Krustal-Wallis test followed by Dunn’s test for post-hoc multiple comparison in Prism 10 software.

### Plaque assays

To analyze the effect of ApMt1 on plaque formation, 200 RH DiCre_T2A Δku80Δhxgprt (parental) or ApMt1^cKD^ tachyzoites per well were used on infected 6-well plates seeded with confluent HFFs and pulsed with 50 nM rapamycin or DMSO (vehicle control) for 2 hours, washed twice with PBS, and replenished with fresh culture medium. The plates were then allowed to grow undisturbed for 8 days before fixing with 100% ethanol and staining with crystal violet. Plaque number and size was quantified using FIJI from two technical replicates for each biological replicate (n=3). Significance was calculated using one-way ANOVA followed by Dunnett’s Test for post hoc multiple comparison in the Prism 10 software.

## Supplemental information

**Table S1**: List of organellar surface proteome candidates

**Table S2**: Apicoplast proteins in *Plasmodium falciparum* with a homolog in *Toxoplasma gondii*

**Table S3**: Expanded list of surface proteome candidates

**Table S4**: List of membrane contact site candidates

**Table S5**: List of plasmids, constructs and primers used in this study

## Notes

### Competing Interest Statement

The authors have declared no competing interest.

